# CrWRKY42 positively regulates chlorophyll degradation and carotenoid biosynthesis in citrus

**DOI:** 10.1101/2023.08.09.552702

**Authors:** Hongyan Chen, Huiyu Ji, Wenkai Huang, Zhehui Zhang, Kaijie Zhu, Shiping Zhu, Lijun Chai, Junli Ye, Xiuxin Deng

## Abstract

Chlorophyll degradation and carotenoid biosynthesis, which occur almost simultaneously during fruit ripening, are essential for coloration and nutritional value of fruits. However, the synergistic regulation of these two processes at transcriptional level remains largely unknown. Here, we identified a novel WRKY transcription factor CrWRKY42 from the transcriptome data of the yellowish bud mutant ‘Jinlegan’ tangor (MT) and its wild type ‘Shiranuhi’ tangor (WT), which was involved in the transcriptional regulation of both chlorophyll degradation and carotenoid biosynthesis pathways. CrWRKY42 activated the expression of *β-carotene hydroxylase 1* (*CrBCH1*) by directly binding to its promoter. Overexpression and interference of *CrWRKY42* in citrus calli demonstrated that *CrWRKY42* promoted carotenoid accumulation by inducing the expression of multiple carotenoid biosynthetic genes. Further assays confirmed that CrWRKY42 also directly bound to and activated the promoters of the genes involved in the carotenoid biosynthesis, including *phytoene desaturase* (*CrPDS*) and *lycopene β-cyclase 2* (*CrLCYB2*). In addition, CrWRKY42 could also bind to the promoter of *STAY-GREEN* (*CrSGR*) and activated its expression, thus promoting chlorophyll degradation. Overexpression and silencing of *CrWRKY42* in citrus fruits indicated that CrWRKY42 positively regulated chlorophyll degradation and carotenoid biosynthesis by synergistically activating the expressions of genes involved in both pathways. In conclusion, our data revealed that CrWRKY42 acted as a positive regulator of chlorophyll degradation and carotenoid biosynthesis to alter the conversion of citrus fruit color. Our findings provide insight into the complex transcriptional regulation of chlorophyll and carotenoid metabolism during fruit ripening.

**One Sentence Summary:** The CrWRKY42 transcription factor coordinates chlorophyll degradation and carotenoid biosynthesis by directly regulating genes involved in these pathways to alter the conversion of citrus fruit color.

## Introduction

Bud sports, an effective way to generate superior cultivars of fruit trees, can alter numerous characteristics, such as fruit texture, size, shape, color, and timing of maturity, of which fruit color mutations are the most common (Jiang et al., 2020). Citrus is the predominant fruit tree in the world and has the highest proportion of sports varieties, with fruit color primarily determined by the combination of chlorophyll and carotenoid (Prasanna et al., 2007). Notably, citrus fruits contain various carotenoids, together with abundant color mutants, providing excellent material for investigating the complex carotenoid metabolism (Zheng et al., 2019).

Genes involved in carotenoid biosynthesis in horticultural plants have been well described (Moise et al., 2014; Nisar et al., 2015). Two geranylgeranyl diphosphate (GGPP) molecules have been reported to be catalyzed by the phytoene synthase (PSY) to synthesize phytoene, and then phytoene is stepwise transformed into lycopene by multiple enzymes such as phytoene desaturase (PDS). The cyclization of lycopene is an important step to divide the major carotenoid synthesis pathway into *α* or *β* branches of the carotenoid biosynthesis pathway (Harjes et al., 2008). In the *β* branch, lycopene is cyclized by lycopene *β*-cyclase (LCYB) to produce *β*-carotene, which is in turn hydroxylated by *β*-carotene hydroxylase (BCH) to successively produce *β*-cryptoxanthin and zeaxanthin (Nisar et al., 2015). The hydroxylation of the BCH-catalyzed *β*-ring of *β*-carotene is essential for carotenoid synthesis. Mutation of *β-carotene hydroxylase* (*CHY2*) leads to the accumulation of *β*-carotene thus producing orange pepper fruits (Borovsky et al., 2013). Silencing *AcBCH1* in kiwifruit results in the increased content of *β*-carotene and the decreased contents of *β*-cryptoxanthin and zeaxanthin (Xia et al., 2022). There are two *β*-carotene hydroxylases BCH1 and BCH2 in citrus, with the hydroxylation capacity and expression level of *BCH1* substantially higher than those of *BCH2* (Zhang et al., 2022b). Silencing *β-carotene hydroxylase* (*Csβ-CHX*) in orange fruit leads to a substantial increase in *β*-carotene, thus producing golden yellow fruits (Pons et al., 2014). The function of *BCH1* has been well characterized in citrus, but the studies on the transcriptional regulation of *BCH1* are limited.

The chlorophyll degradation pathway has been well elucidated in plants (Biswal et al., 2013). Chlorophyll b is converted to chlorophyll a by the NON-YELLOW COLORING (NYC) and subsequently catabolized by different dephytylation enzymes such as STAY-GREEN (SGR), pheophytin pheophorbide hydrolase (PPH), and Pheophorbide a monooxygenase (PAO) (Christ et al., 2014; Kuai et al., 2018; Jiao et al., 2020). Notably, SGR encodes magnesium-dechelatase, which is known as Mendel’s green cotyledon gene (Shimoda et al., 2016). SGR has been identified as a key regulator of chlorophyll degradation in numerous species, including *Arabidopsis* (Sakuraba et al., 2012), tomato (Barry et al., 2008), and litchi (Zou et al., 2023). In citrus, SGR can coordinate chlorophyll degradation with carotenoid biosynthesis during fruit ripening (Zhu et al., 2021b). However, studies on the transcriptional regulation of SGR are still limited.

In recent years, numerous transcription factors (TFs) have been reported to be involved in plant growth and development. WRKY TFs, one of the most essential families of plant-specific TFs, are extensively involved in plant growth and development by recognizing the cis-acting element W-box (Pandey and Somssich, 2009). WRKY TFs function in multiple biological processes such as seed maturation and dormancy (Ding et al., 2014), biotic and abiotic stresses response (Tang et al., 2022; Wang et al., 2022), cell development (Tang et al., 2023), senescence (Zhao et al., 2020), secondary metabolite synthesis and regulation, and fruit ripening (Schluttenhofer and Yuan, 2015). FvWRKY48 controls fruit softening and ripening by binding to the pectate lyase *FvPLA* promoter in *Fragaria vesca* (Zhang et al., 2022a). In *Sorghum bicolor* L., SbWRKY50 negatively regulates chlorophyll degradation by directly binding to the promoters of several chlorophyll catabolic genes (Chen et al., 2023b). In addition, OfWRKY3 binds the W-box palindrome motif in the carotenoid cleavage dioxygenase gene *OfCCD4* promoter to induce its expression in *Osmanthus fragrans* (Han et al., 2016). SlWRKY35 positively regulates carotenoid biosynthesis by activating the expression of the deoxy-D-xylulose 5-phosphate synthase (*SlDXS1*) gene in tomato fruit (Yuan et al., 2022). However, few reports on the roles of WRKY TFs in the coordinated regulation of carotenoid biosynthesis and chlorophyll degradation have been available.

‘Jinlegan’ (MT) tangor is a natural mutant of ‘Shiranuhi’ (WT) tangor (*Citrus reticulata*), whose fruit has bright yellow color. In the previous study, we have found an increased content of *β*-carotene and a decreased content of *β*-cryptoxanthin and zeaxanthin in the flavedo of ‘Jinlegan’ (MT) tangor compared to the wild type ‘Shiranuhi’ (WT) tangor at the breaker stage (190 DAF), which might be due to the reduced expression level of *CrBCH1* (Chen et al., 2023a). However, the underlying mechanism of regulation are not clear. In this study, we identified a transcription factor *CrWRKY42* from the transcriptome data of the yellowish bud mutant ‘Jinlegan’ tangor and ‘Shiranuhi’ tangor, and this TF was involved in the regulation of carotenoids biosynthesis and conversion of citrus fruits color. We found that CrWRKY42 activated the expression of carotenoid biosynthesis genes *CrBCH1* as well as *CrPDS* and *CrLCYB2* by directly binding to their promoters, thus promoting the accumulation of carotenoid. In addition, CrWRKY42 also directly bound to chlorophyll degradation gene *CrSGR* and activated its expression to promote the degreening process. In summary, this study indicated that CrWRKY42 acted as a positive regulator of chlorophyll degradation and carotenoid biosynthesis to accelerate the conversion of citrus fruit color. Our findings provide insight into the complex transcriptional regulation of plant chlorophyll and carotenoid metabolism.

## Results

### CrWRKY42 is a nucleus-localized transcriptional activator

Sequence analysis revealed that CrWRKY42 was a transcription factor with a WRKY domain and belonged to the group Ⅲ subfamily (Figure.1A). We found *CrWRKY42* was significantly down-regulated along with *CrBCH1* at the breaker stage (190 DAF) in MT (Figure. 1B, C), suggesting that *CrWRKY42* might regulate the expression of *CrBCH1* to affect carotenoid biosynthesis. Subcellular localization experiments indicated that CrWRKY42 was localized to the nucleus (Figure. 2A).

**Figure 1.**
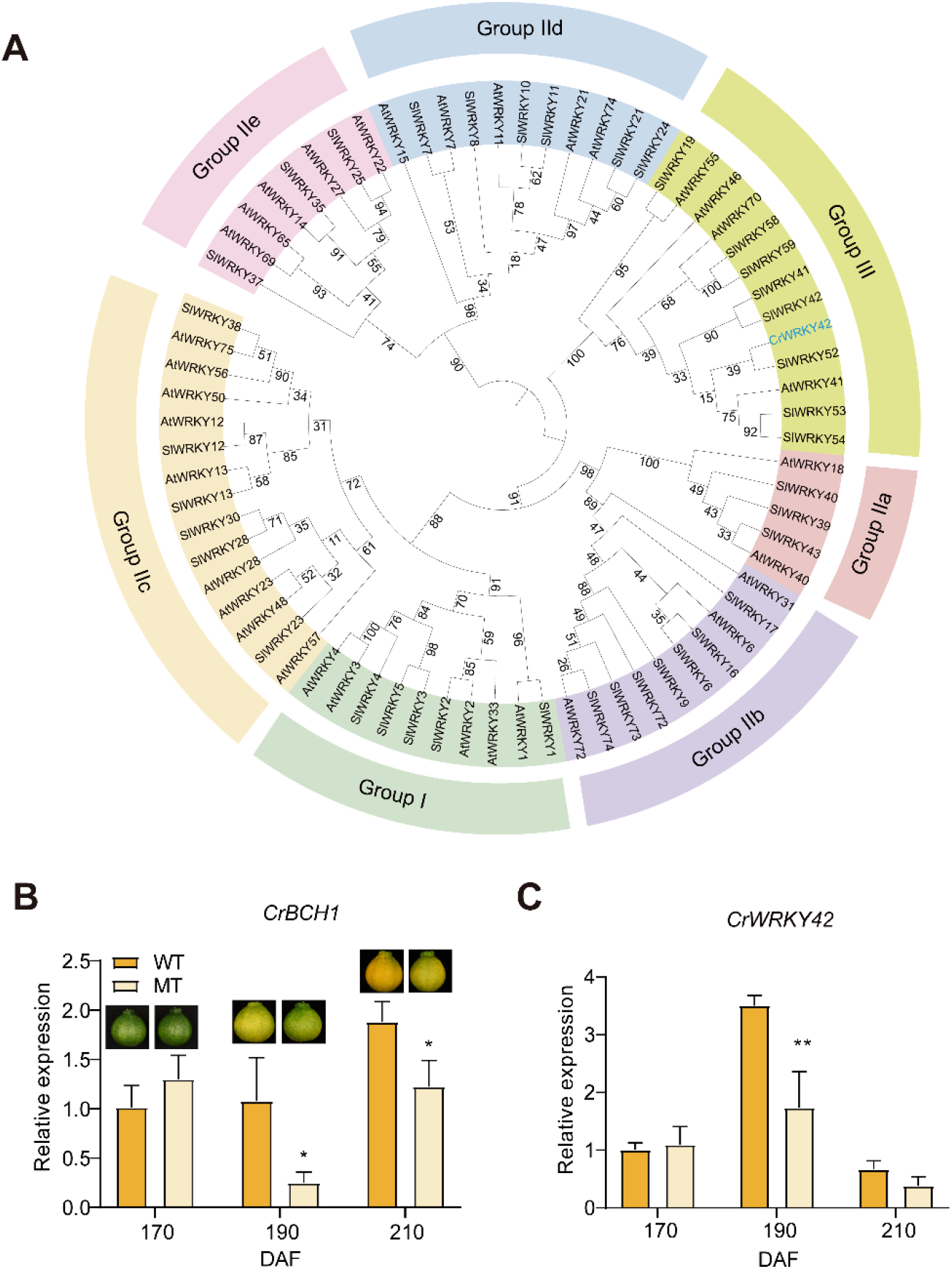
CrWRKY42 acts as a candidate regulatory gene for carotenoid biosynthesis. (A) Phylogenetic trees of CrWRKY42 and partial WRKY genes in Arabidopsis and tomato. (B) Relative expression of *CrBCH1* in WT (‘Shiranuhi’) and MT (‘Jinlegan’) flavedo at different developmental stages. (C) Relative expression of *CrWRKY42* in WT and MT flavedo at different developmental stages. (Student’s *t*-test, * *P*<0.05, ** *P*<0.01).

**Figure 2.**
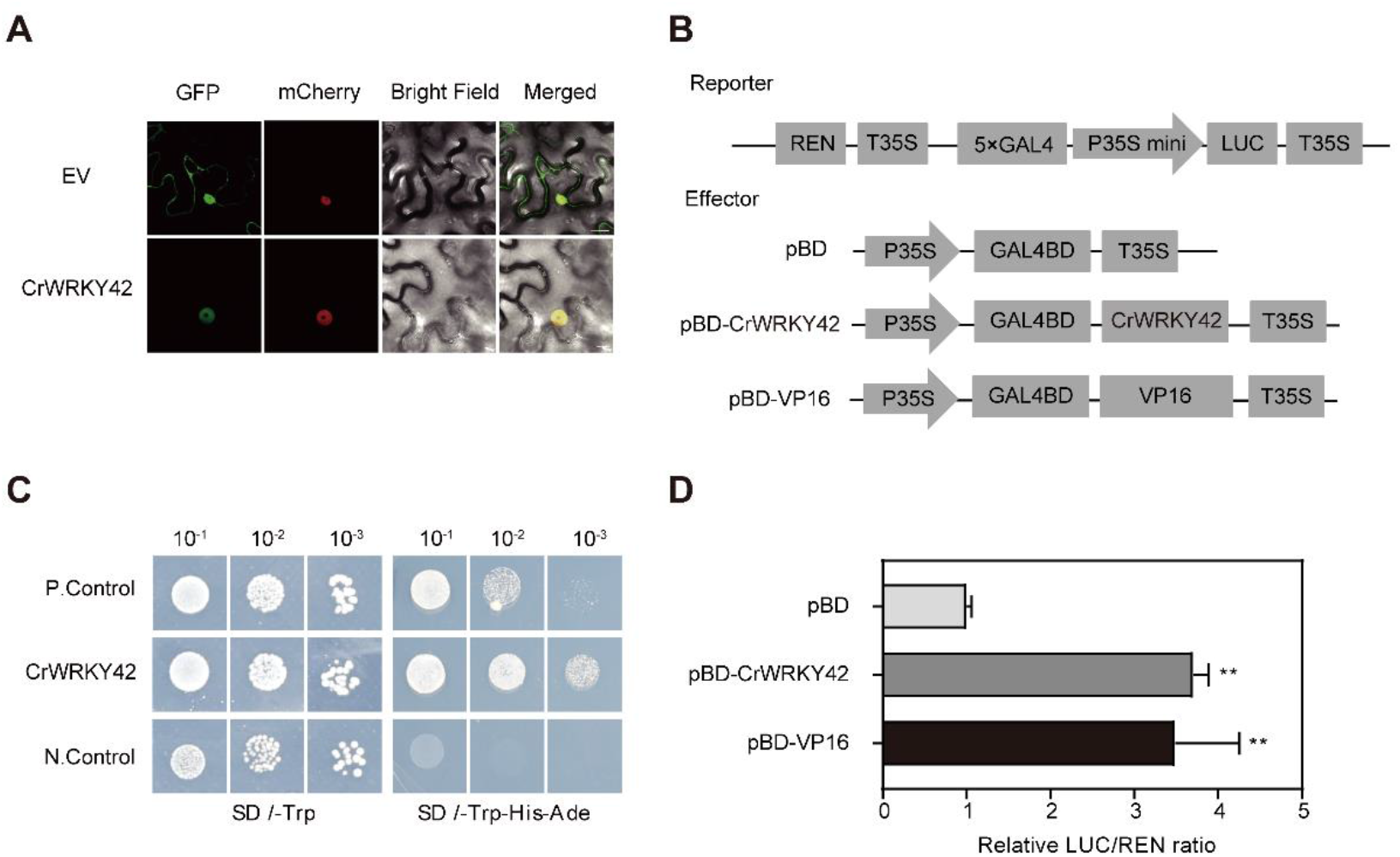
Subcellular localization and transcriptional activity of CrWRKY42. (A) Subcellular localization of CrWRKY42. CrWRKY42-GFP fusion vector and nuclear marker H2B were transiently transformed into *Nicotiana benthamiana* leaves, and localization was observed under confocal laser microscopy. EV, empty vector pRI121; CrWRKY42, vector pRI121 containing CrWRKY42; GFP, greenfluorescence; mCherry, redfluorescence; Bright Field, white light; Merged, combined GFP and mCherry signals. Bar=100 μm. (B) Schematic diagram of the dual luciferase vector. pBD, negative control; CrWRKY42, vector pBD containing CrWRKY42; pBD-VP16, positive control. (C) Transactivation analysis of CrWRKY42 using yeast assay. CrWRKY42, vector pGBKT7 containing CrWRKY42; P.Control (pGBKT7-53 + pGADT7-RecT) and N.Control (empty vector pGBKT7) served as the positive and negative controls, respectively. The yeast cells containing the above-mentioned vectors, respectively, were grown on SD/-Trp or SD/-Trp-His-Ade medium at graded dilution concentrations for transcriptional activity detection. (D) Relative transcriptional activity of CrWRKY42 using dual luciferase assay. The data were expressed as mean ± standard deviation (SD) of five biological replicates. Asterisks indicate statistically significant differences (one-way ANOVA, ** *P*<0.01).

To investigate the transcriptional activity of CrWRKY42 *in vivo*, a fusion vector of pGBKT7-CrWRKY42 was constructed and transformed into yeast (*Saccharomyces cerevisiae*) strain AH109. The results demonstrated that only yeast cells carrying positive control and pGBKT7-CrWRKY42 survived on SD/-Trp-His-Ade medium, while the negative control did not survive, indicating that CrWRKY42 had transcriptional activation activity (Figure. 2C). Further, we investigated the transcriptional activity of CrWRKY42 using dual luciferase system. Compared with the negative control, CrWRKY42 significantly enhanced the activity of the luciferase (LUC) reporter, which was comparable to the level of the positive control pBD-VP16 (Figure. 2B, D). These results confirmed that CrWRKY42 acted as a transcriptional activator.

### CrWRKY42 directly binds and activates the promoter of *CrBCH1*

We hypothesized that CrWRKY42 might act as an upstream regulator of *CrBCH1*. To test our hypothesis, yeast one-hybrid assay (Y1H), electrophoretic mobility shift assay (EMSA), and dual luciferase assay were performed to examine whether CrWRKY42 bound to the *CrBCH1* promoter.

The results showed that yeast cells carrying the CrWRKY42 (pGADT7-CrWRKY42 + pAbAi-*CrBCH1pro*) and positive control (pGADT7-Recp53 + p53-AbAi) survived on the selective medium supplemented with 125 ng mL^-1^ aureobasidin A (AbA), while the negative control (pGADT7 + pAbAi-*CrBCH1pro*) did not (Figure. 3A), indicating that CrWRKY42 interacted with the promoter of *CrBCH1* in the yeast system. In addition, the ability of CrWRKY42 to bind the W-box on *CrBCH1* promoter was further investigated by EMSA. The EMSA results showed that shifted band was observed when the recombinant protein MBP-CrWRKY42 and a labeled probe co-existed, and that this shifted band was weakened by the unlabeled probe, whereas it was not weakened by the mutated unlabeled probe (Figure. 3D). These results suggested that the CrWRKY42 protein specifically bound to the W-box on *CrBCH1* promoter *in vitro*.

**Figure 3.**
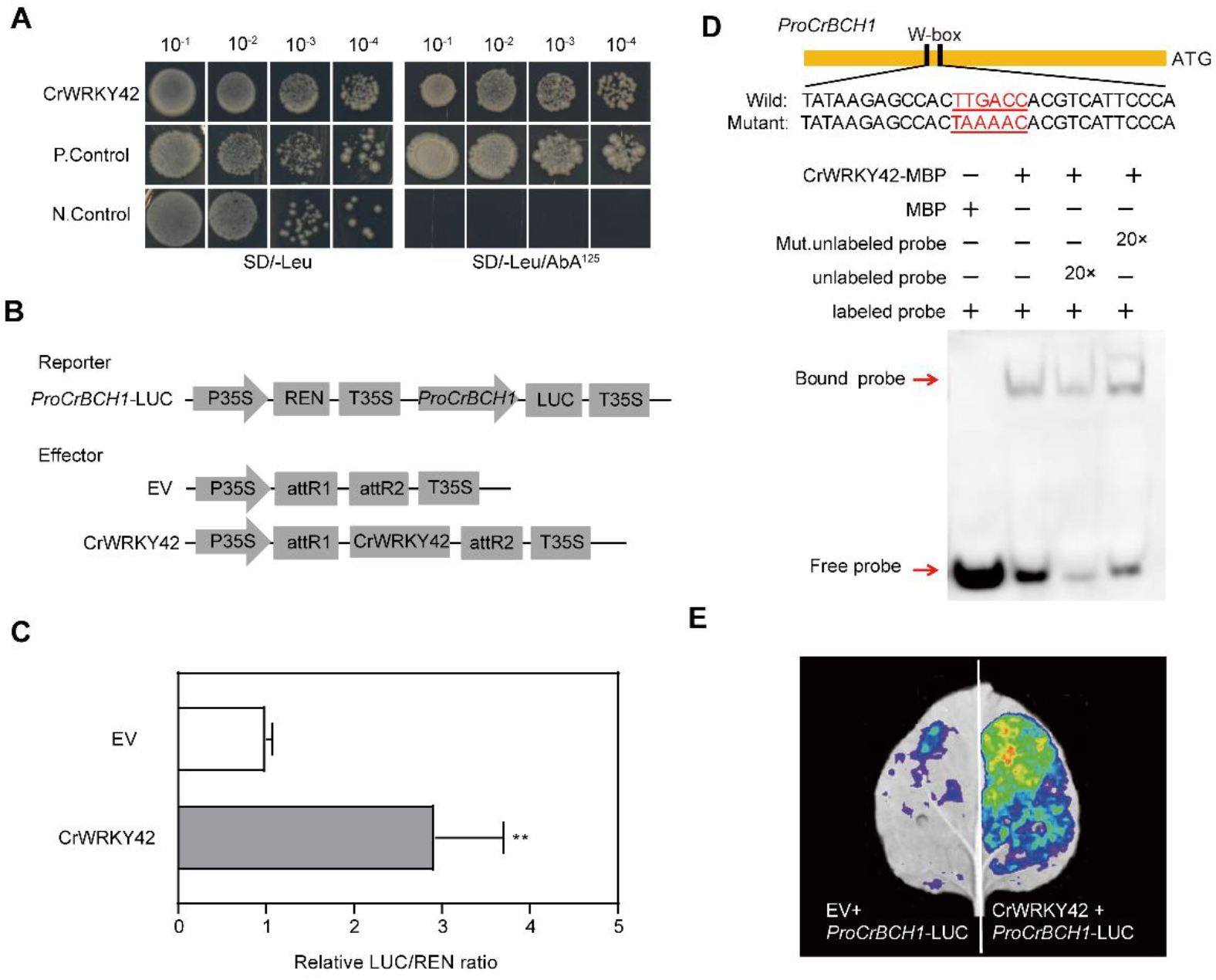
CrWRKY42 directly binds to the promoter of *CrBCH1* and activates its expression. (A) Yeast one-hybrid assay (Y1H) showing the binding of CrWRKY42 to the promoter of *CrBCH1*. CrWRKY42 indicated yeast cells carrying pGADT7-CrWRKY42 + pAbAi-*CrBCH1pro*. P.Control (pGADT7-Recp53 + p53-AbAi) and N.Control (pGADT7 + pAbAi-*CrBCH1pro*) served as the positive and negative controls, respectively. SD/-Leu/AbA^125^ was SD/-Leu medium supplemented with 125 ng ml^-1^ aureobasidin A. (B) Schematic diagram of vectors used for the dual luciferase assay. EV, empty vector PK7WG2D; CrWRKY42, vector PK7WG2D containing CrWRKY42. (C) Dual luciferase assay showing relative CrWRKY42 activation to the promoter of *CrBCH1*. (D) Binding of CrWRKY42 protein to the W-box on the promoter of *CrBCH1.* Electrophoretic mobility shift assay (EMSA) was performed with a biotin-labeled *CrBCH1* promoter fragment containing W-box (TTGACC). Unlabeled same *CrBCH1* promoter fragment and mutated motifs (TAAAAC) were used as competitors. MBP protein was used as a negative control. ‘+’ and ‘−’ indicate the presence and absence of the specified probe or protein, respectively. (E) Dual luciferase imaging demonstrating relative CrWRKY42 activation to the promoter of *CrBCH1*.The data were expressed as mean ± standard deviation (SD) of five biological replicates. Asterisks indicate statistically significant differences (Student’s *t* test, ** *P*<0.01).

To further determine whether CrWRKY42 could activate the expression of *CrBCH1*, we performed dual luciferase assay. The promoter of *CrBCH1* was inserted into the LUC reporter vector, and the CDS of CrWRKY42 was cloned into the effector vector (Figure 3B). The results showed that compared with empty vector, CrWRKY42 significantly elevated the relative luciferase expression (LUC/REN ratio) driven by the *CrBCH1* promoter (Figure. 3C). Furthermore, luciferase imaging assays also confirmed that CrWRKY42 activated the *CrBCH1* promoter. Coexpression region (CrWRKY42 + *CrBCH1pro*-Luc) exhibited a strong fluorescent signal, whereas the control region (EV + *CrBCH1pro*-Luc) displayed a weak fluorescent signal (Figure. 3E), indicating that CrWRKY42 activated the *CrBCH1* promoter *in vivo*. Taken together, these results above indicated that CrWRKY42 protein directly bound to the promoter of *CrBCH1* and activated its expression.

### CrWRKY42 alters carotenoid biosynthesis by affecting expression of key carotenoid pathway genes

To investigate the role of CrWRKY42 in carotenoid biosynthesis in citrus, we overexpressed it in citrus calli. The *CrWRKY42*-overexpression lines exhibited stable phenotype with more yellow color than the control (WT) (Figure. 4A). Semi-quantitative PCR and RT-qPCR results showed that *CrWRKY42* had higher expression level in three overexpression lines (OE-2, OE-3, and OE-4) than in the control (Figure. 4B, C), and thus these three overexpression lines were subjected to further phenotypic analysis. Consistent with the phenotypic changes, the total carotenoid content was significantly higher in *CrWRKY42*-overexpression lines than in the control (Figure. 4D). The contents of zeaxanthin and violaxanthin significantly increased, compared with those in the control, but no significant difference in lutein content was observed between the *CrWRKY42*-overexpression lines and the control. Notably, *β*-carotene content exhibited a significant decrease in *CrWRKY42*-overexpression lines, compared with that in the control (Figure. 4D).

**Figure 4.**
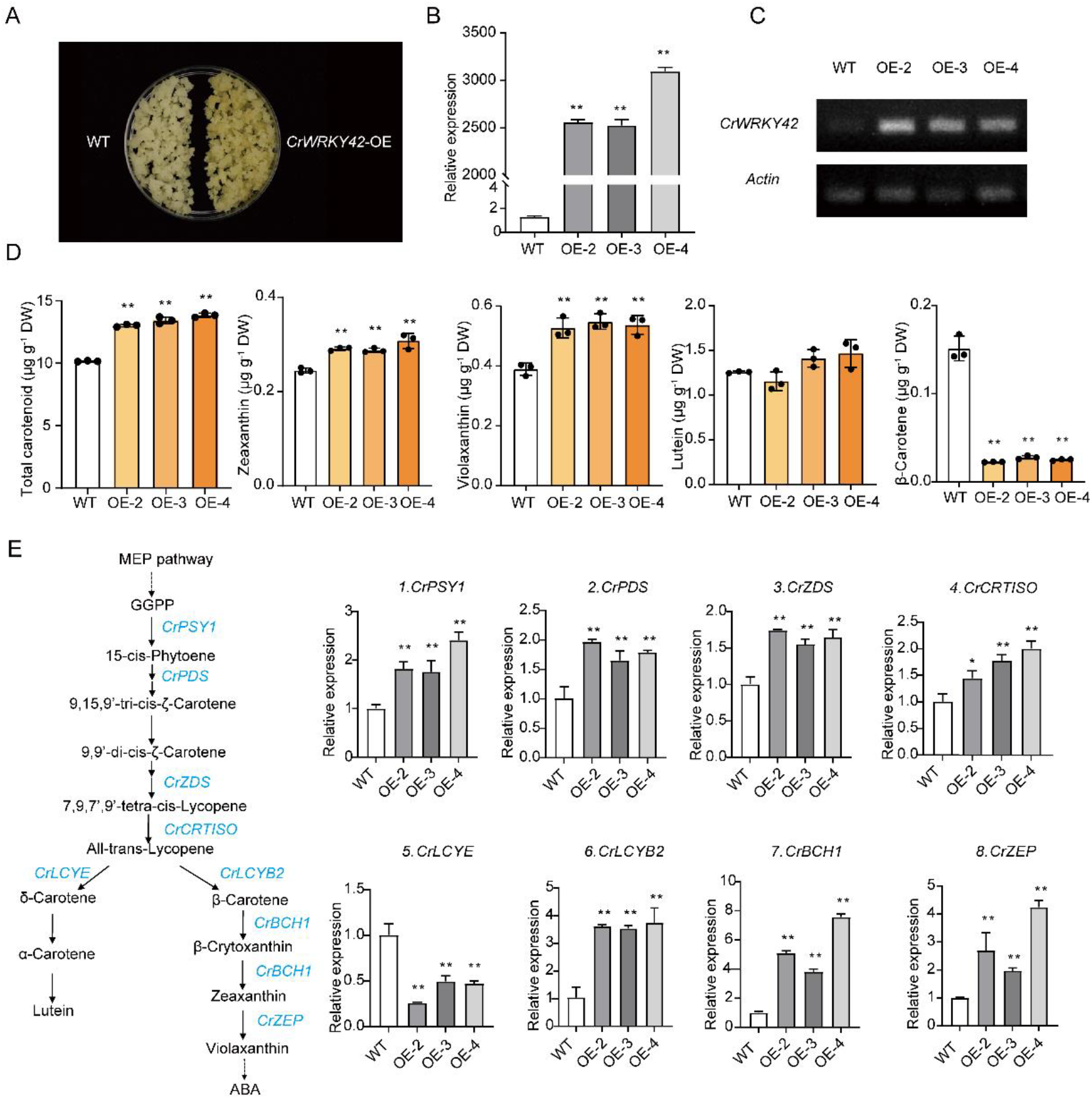
Overexpression of *CrWRKY42* promotes carotenoid accumulation in citrus calli. (A) Phenotypes of *CrWRKY42*-overexpression citrus calli. WT, wild type of Marsh grapefruit. (B) Expression of *CrWRKY42* in transgenic citrus calli. OE-2, OE-3, and OE-4 indicated three independent *CrWRKY42*-overexpression citrus calli lines. (C) Semi-quantitative RT-PCR analysis of *CrWRKY42* transcript levels. *Actin* was used as an internal reference. (D) Carotenoid content in *CrWRKY42*-overexpression lines. (E) Expression of carotenoid biosynthesis genes in *CrWRKY42*-overexpression lines. *CrPSY1*, phytoene synthase 1; *CrPDS*, phytoene desaturase; *CrZDS*, *ζ*-carotene desaturase; *CrCRTISO*, carotene isomerase; *CrLCYE*, lycopene *ε*-cyclase; *CrLCYB2*, lycopene *β*-cyclase 2; *CrBCH1*, *β*-carotene hydroxylases 1; *CrZEP*, zeaxanthin epoxidase. DW, dry weight. The data were expressed as mean ± standard deviation (SD) of three biological replicates. Asterisks indicate statistically significant differences (Student’s *t* test, * *P*<0.05, ** *P*<0.01).

To further explore the underlying mechanism of carotenoid content variations in *CrWRKY42*-overexpression lines, we examined the expression levels of key carotenoid biosynthesis genes by RT-qPCR. The results showed that the expressions of most genes involved in carotenoid biosynthesis including *CrPSY1*, *CrPDS*, ζ-carotene desaturase (*CrZDS*), carotene isomerase (*CrCRTISO*), *CrLCYB2*, *CrBCH1*, and zeaxanthin epoxidase (*CrZEP*) were remarkably elevated, while the expression of lycopene *ε*-cyclase (*CrLCYE*) was lowered (Figure. 4E). It should be noted that the expression of *CrBCH1* was elevated by 4-7 folds, which might be responsible for the reduction of *β*-carotene in the *CrWRKY42*-overexpression lines. **Interference of *CrWRKY42* expression in citrus calli reduces carotenoid accumulation**

*CrWRKY42* expression was further interfered with in citrus calli to examine the transcriptional regulatory role of CrWRKY42 in carotenoid biosynthesis. In contrast to the *CrWRKY42*-overexpression lines, *CrWRKY42*-RNAi lines displayed a slightly paler color (Figure. 5A). The expression level of *CrWRKY42* was significantly lower in *CrWRKY42*-RNAi lines (RNAi-1, RNAi-2, RNAi-3) than in the control (WT) (Figure. 5C). Compared to that in the control, the total carotenoid content in *CrWRKY42*-RNAi lines was significantly decreased (Figure. 5B), which was consistent with the reduced expression of the carotenoid biosynthesis genes *CrPSY1*, *CrPDS*, *CrZDS*, *CrCRTISO*, *CrZEP*, and carotenoid cleavage dioxygenase 1 (*CrCCD1*) (Figure. 5C).

**Figure 5.**
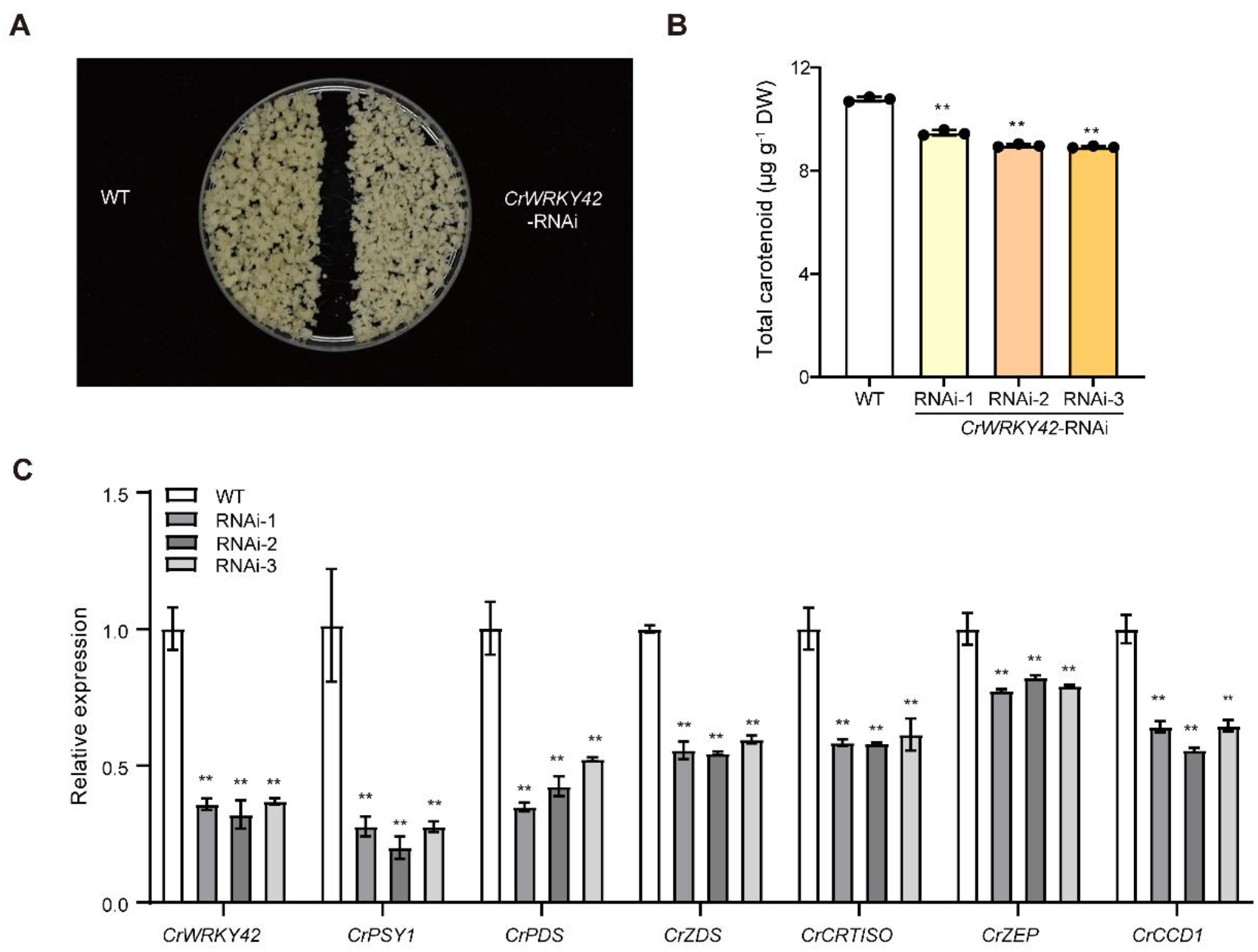
Interference of *CrWRKY42* reduces carotenoid accumulation in citrus calli. (A) Phenotype of *CrWRKY42*-RNAi citrus calli. WT, wild type of Marsh grapefruit. (B) Total carotenoid content in *CrWRKY42*-RNAi citrus calli. RNAi-1, RNAi-2, RNAi-3, three independent *CrWRKY42* interference lines. (C) Expression levels of *CrWRKY42* and carotenoid biosynthesis genes in *CrWRKY42*-RNAi lines. *CrPSY1*, phytoene synthase 1; *CrPDS*, phytoene desaturase; *CrZDS*, *ζ*-carotene desaturase; *CrCRTISO*, carotene isomerase; *CrZEP*, zeaxanthin epoxidase. *CrCCD1,* carotenoid cleavage dioxygenase 1. DW, dry weight. The data were expressed as mean ± standard deviation (SD) of three biological replicates. Asterisks indicate statistically significant differences (Student’s *t* test, ** *P*<0.01).

### CrWRKY42 positively regulates chlorophyll degradation and carotenoid accumulation in citrus fruit

Due to the long juvenile period and complex genetic background of citrus, obtaining transgenic citrus fruits is challenging and time-consuming, thus the transient transformation is a promising method to investigate the function of genes in citrus (Gong *et al*., 2021*b*). The transient transformation has also been proved to be effective in various fruits, such as tomato, apple, and strawberry (Hoffmann *et al*., 2006; Orzaez *et al*., 2006; An *et al*., 2018).

To better investigate the function of CrWRKY42 in citrus fruit, we performed transient overexpression and interference experiments. *Agrobacterium tumefaciens* carrying *CrWRKY42* and empty vector (PK7-EV) were injected into the peel of citrus fruits and stored at room temperature for 12 days, respectively. We observed that the peel color near the injection site of *CrWRKY42* turned yellow in overexpression lines, while that near the injection site of PK7-EV was green in control fruit (Figure. 6A), which was supported by a significantly higher citrus color index (CCI) near the *CrWRKY42* injection site than the control (Figure. 6C). Strong fluorescence signal of GFP was observed on the inner side of the peel (Figure. 6B). The expression level of *CrWRKY42* significantly elevated in citrus peel of *CrWRKY42*-overexpression fruits, compared to that in the control, indicating that *CrWRKY42* was successfully overexpressed (Figure. 6E). Compared with that in the control, total carotenoid content was considerably increased in *CrWRKY42*-overexpression fruits (Figure. 6D), while chlorophyll content was decreased (Figure. 6F). Consistently, the expression of *CrDXS*, *CrPSY1*, *CrZDS*, *CrCRTISO*, and *CrBCH1* related to carotenoid biosynthesis were significantly increased (Figure. 6E). In addition, we observed that chlorophyll degradation was faster in the *CrWRKY42*-overexpression lines than in the control. Therefore, we further examined the expression level of genes related to chlorophyll degradation and found that *CrNYC*, *CrSGR*, *CrPPH*, and *CrPAO* were significantly up-regulated in the *CrWRKY42*-overexpression lines (Figure. 6G). These results jointly suggested that CrWRKY42 positively regulated chlorophyll degradation and carotenoid accumulation, thus promoting citrus fruit color conversion.

**Figure 6.**
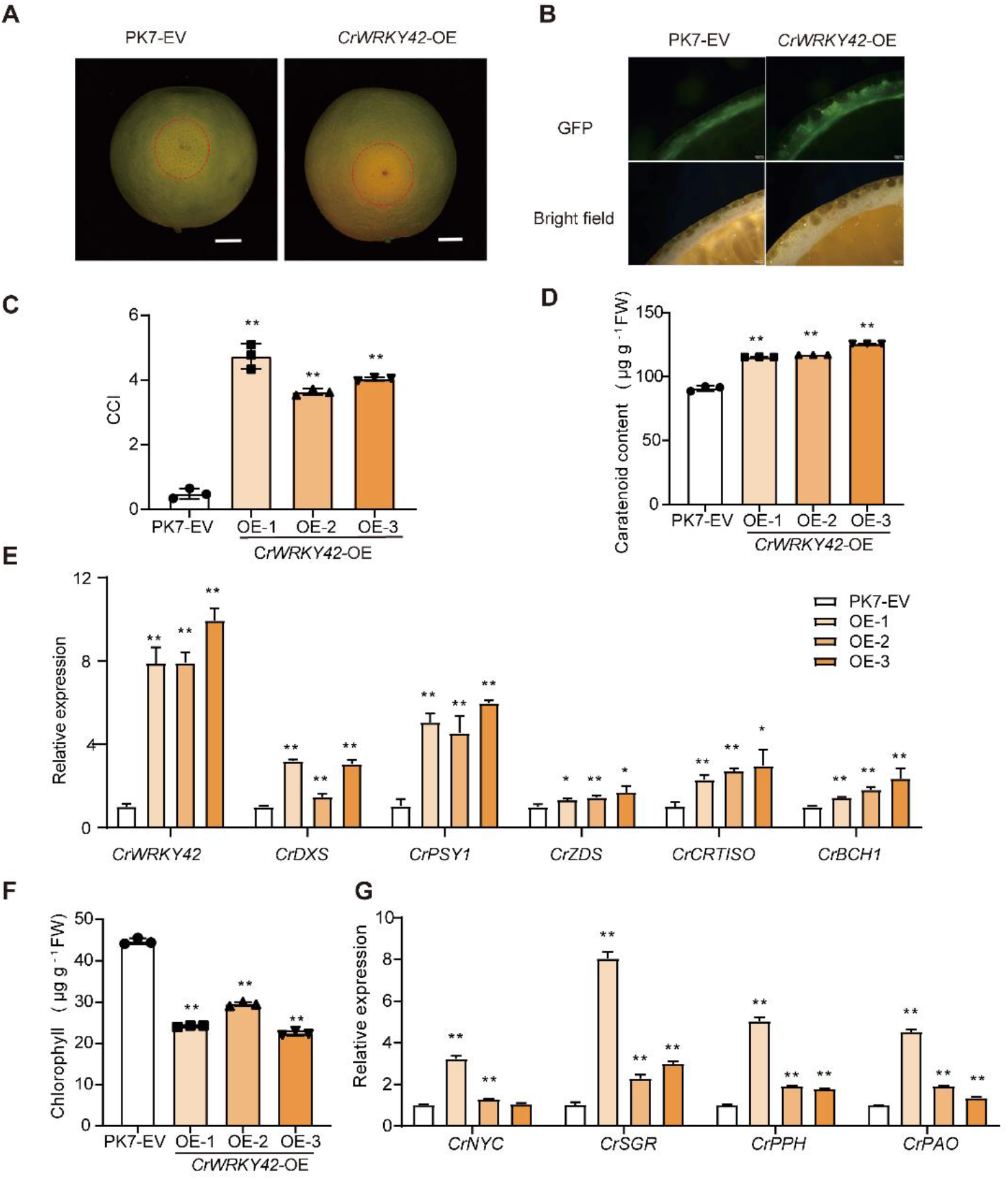
Transient overexpression of *CrWRKY42* promotes chlorophyll degradation, and carotenoid accumulation, and color conversion in citrus fruits. (A) Phenotype of *CrWRKY42-*overexpression citrus fruits around the injection site. PK7-EV indicated empty vector used as negative control. Bar = 1cm. (B) GFP fluorescence observation of *CrWRKY42-*overexpression citrus fruits. Bar = 1mm. (C) CCI values of *CrWRKY42-*overexpression citrus fruits around the injection site. (D) Total carotenoid content around the injection sites in *CrWRKY42-*overexpression citrus fruits. (E) Expression levels of *CrWRKY42* and carotenoid biosynthesis genes in *CrWRKY42-*overexpression citrus fruits. (F) Total chlorophyll content around the injection sites in *CrWRKY42-*overexpression citrus fruits. (G) Expression levels of chlorophyll degradation genes in *CrWRKY42-*overexpression citrus fruits. CCI, citrus color index; *CrDXS*, deoxy-D-xylulose 5-phosphate synthase; *CrPSY1*, phytoene synthase 1; *CrZDS*, *ζ*-carotene desaturase; *CrCRTISO*, carotene isomerase; *CrBCH1*, *β*-carotene hydroxylases 1; *CrNYC*, NONYELLOW COLORING; *CrSGR*, STAY-GREEN; *CrPPH*, pheophytinase; *CrPAO*, pheophorbide a monooxygenase; FW, fresh weight. The data were expressed as mean ± standard deviation (SD) of three biological replicates. Asterisks indicate statistically significant differences (Student’s *t* test, * *P*<0.05, \*\**P*<0.01).

In addition, we constructed the *CrWRKY4*2-RNAi vector and injected it into the peel of citrus fruits at the breaker stage, and citrus fruits injected with an empty vector (RNAi-EV) were used as negative control. After 18 days of dark treatment, the color of the peel near the *CrWRKY42-*RNAi injection site remained a light green, while the color near the injection site in the control completely turned yellow (Figure. 7A), indicating that *CrWRKY42* interference can delay degreening. Strong fluorescence signal of GFP was also observed at the inner side of the peel (Figure. 7B), and the expression level of *CrWRKY42* was significantly down-regulated in the *CrWRKY42-*RNAi fruits (Figure. 7F). Consistently, the value of CCI and total carotenoid content near the injection site of *CrWRKY42-*RNAi were significantly lower than those in the control (Figure. 7C, D). Total chlorophyll content was higher in *CrWRKY42-*RNAi fruits than in the control (Figure. 7E). Additionally, the expressions of carotenoid biosynthesis genes *CrZDS*, *CrBCH1*, *CrLCYB2*, *CrNCED2* and the chlorophyll degradation gene *CrSGR* also displayed significant down-regulation in the *CrWRKY42-*RNAi lines (Figure. 7F). Taken together, CrWRKY42 positively regulated citrus fruit coloration by simultaneously modulating the expression of carotenoid biosynthesis genes and chlorophyll degradation genes to promote carotenoid accumulation and chlorophyll degradation.

**Figure 7.**
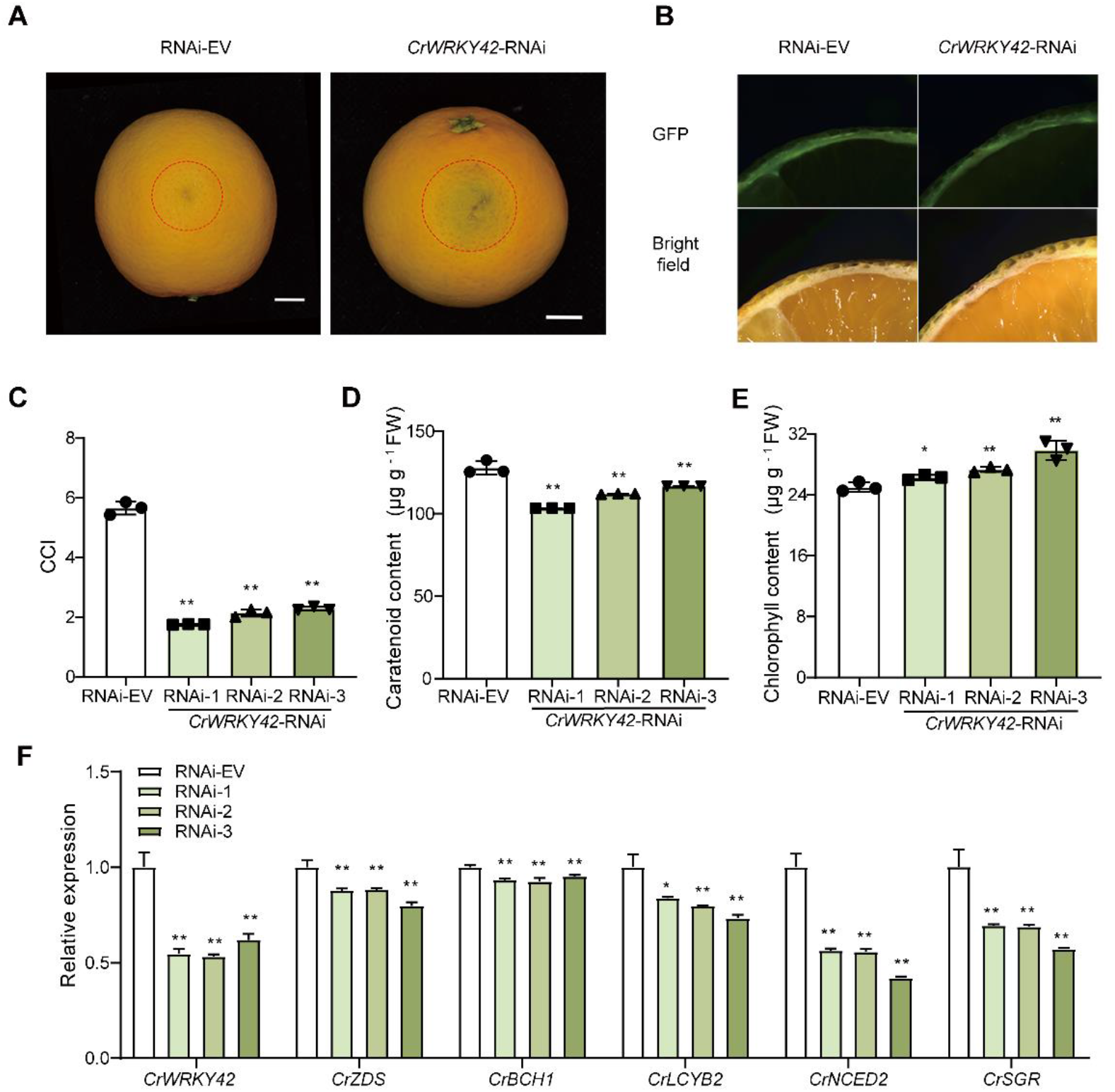
Transient interference of *CrWRKY42* delays color conversion in citrus fruits. (A) Phenotype of *CrWRKY42-*RNAi citrus fruits. RNAi-EV indicated the RNAi empty vector used as negative control. *CrWRKY42-*RNAi denoted RNAi vector containing CrWRKY42. Bar = 1cm. (B) GFP fluorescence observation of *CrWRKY42-*RNAi citrus fruits. Bar = 1mm. (C) CCI values of *CrWRKY42-*RNAi citrus fruits. (D) Total carotenoid content around the injection sites in *CrWRKY42-*RNAi citrus fruits. (E) Total chlorophyll content around the injection sites in *CrWRKY42-*RNAi citrus fruits. (F) Expression levels of *CrWRKY42*, carotenoid biosynthesis genes, and chlorophyll degradation genes in *CrWRKY42-*RNAi citrus fruits. CCI, citrus color index; *CrZDS*, *ζ*-carotene desaturase; *CrBCH1*, *β*-carotene hydroxylases 1; *CrLCYB2*, lycopene *β*-cyclase 2; *CrNCED2*, 9-cis-epoxycarotenoid dioxygenase 2; *CrSGR*, STAY-GREEN; FW, fresh weight. The data were expressed as mean ± standard deviation (SD) of three biological replicates. Asterisks indicate statistically significant differences (Student’s *t* test, * *P*<0.05, ** *P*<0.01).

### CrWRKY42 directly binds to and activates the promoter of *CrSGR, CrPDS*, and *CrLCYB2*

In both *CrWRKY42*-overexpression and *CrWRKY42-*RNAi fruits, we observed the alterations of chlorophyll degradation consistent with the expression of *CrSGR*, and thus we hypothesized that CrWRKY42 might be an upstream regulator of *CrSGR*. To test our hypothesis, we performed EMSA, Y1H, and dual luciferase assay. EMSA results demonstrated that CrWRKY42 bound to the W-box on the promoter of *CrSGR* (Figure. 8A). Furthermore, CrWRKY42 interacted with *CrSGR* promoter in the yeast system (Figure. 8D). Dual luciferase assay indicated that compared to empty vector, CrWRKY42 exhibited a higher capacity to significantly activate the expression of *CrSGR* promoter (Figure. 8E). Taken together, these results suggested that CrWRKY42 could directly bind to the W-box on the promoter of *CrSGR* and activate its expression.

**Figure 8.**
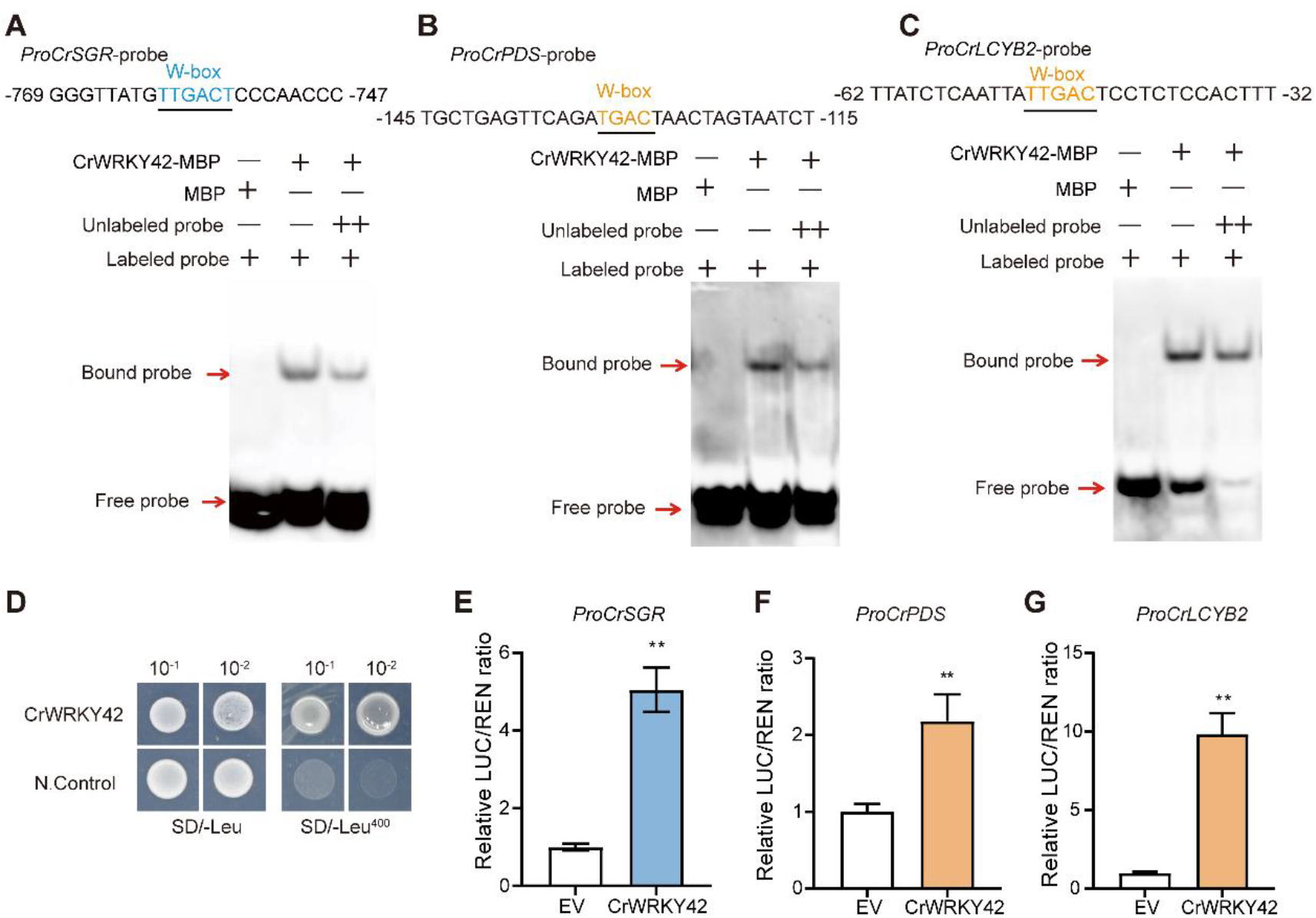
Chlorophyll degradation and carotenoid biosynthesis genes are activated by CrWRKY42. (A) Binding of CrWRKY42 protein to the W-box on the promoter of *CrSGR* using electrophoretic mobility shift assay (EMSA). EMSA was performed with a biotin-labeled *CrSGR* promoter fragment containing W-box (TTGACT). Unlabeled same *CrSGR* promoter fragment was used as competitor. MBP protein was used as a negative control. ‘+’ and ‘−’ indicate the presence and absence of the specified probe or protein, respectively. (B) Binding of CrWRKY42 protein to the W-box on the promoter of *CrPDS* using EMSA. (C) Binding of CrWRKY42 protein to the W-box on the promoter of *CrLCYB2* using EMSA. (D) Binding of CrWRKY42 to the promoter of *CrSGR* using yeast one-hybrid assay (Y1H). CrWRKY42, (pGADT7-CrWRKY42 + pAbAi-*CrSGRpro*). N.Control (pGADT7 + pAbAi-*CrBCH1pro*) served as the negative control. SD/-Leu/AbA^400^ indicated SD/-Leu medium supplemented with 400 ng ml^-1^ aureobasidin A. (E) Dual luciferase assay showing relative CrWRKY42 activation to the promoter of *CrSGR.* (F) Dual luciferase assay showing relative CrWRKY42 activation to the promoter of *CrPDS*. (G) Dual luciferase assay showing relative CrWRKY42 activation to the promoter of *CrLCYB2*. *CrSGR*, STAY-GREEN. *CrPDS*, phytoene desaturase; *CrLCYB2,* lycopene *β*-cyclase 2; The data were expressed as mean ± standard deviation (SD) of five biological replicates. Asterisks indicate statistically significant differences (Student’s *t* test, ** *P*<0.01).

In addition, numerous genes in the carotenoid biosynthesis pathway were up-regulated in both *CrWRKY42*-overexpression citrus calli and fruits, based on which, we speculated that these genes might also serve as potential target genes of CrWRKY42. EMSA results demonstrated that CrWRKY42 bound to the W-box on the promoter of *CrPDS* and *CrLCYB2* (Figure. 8B, C). The results of dual luciferase assay indicated that compared to the control (empty vector), CrWRKY42 significantly activated the expressions of promoters of *CrPDS* and *CrLCYB2* (Figure. 8F, G). In conclusion, in addition to activating the promoter of *CrBCH1*, CrWRKY42 directly bound to and activated the promoters of other essential carotenoid biosynthesis genes *CrPDS* and *CrLCYB2*, as well as the essential chlorophyll degradation gene *C*r*SGR*.

## Discussion

Chlorophyll degradation and carotenoid biosynthesis normally occur simultaneously during fruit ripening, which is essential for vibrant fruit coloration. This study found that these two pathways were co-regulated by CrWRKY42 in citrus. CrWRKY42 coordinates chlorophyll degradation and carotenoid biosynthesis by directly regulating genes involved in these two pathways to alter the conversion of citrus fruit color (Figure 9). Our study provides insight into the complex transcriptional regulation of chlorophyll degradation and carotenoid biosynthesis during fruit ripening.

**Figure 9.**
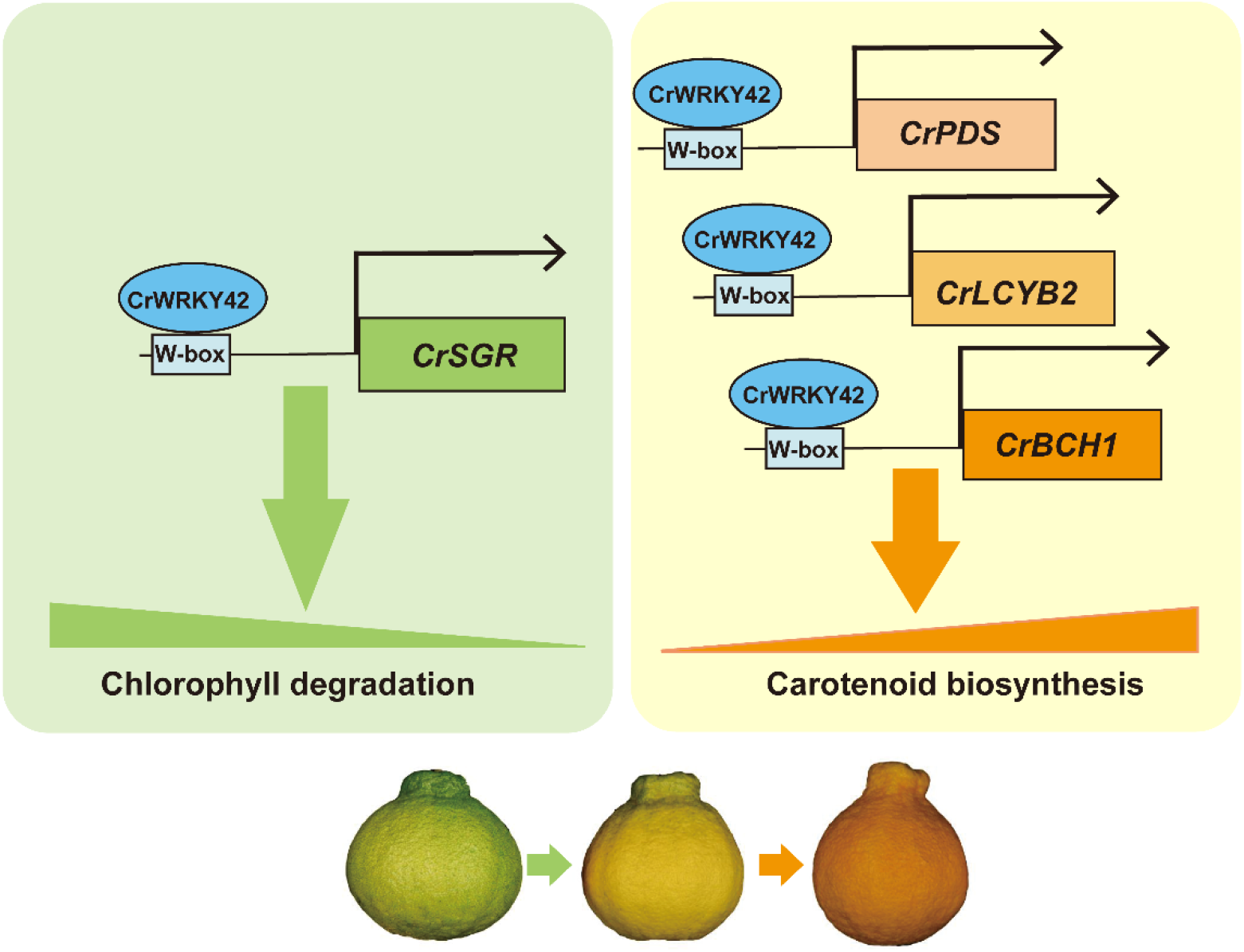
Model diagram of CrWRKY42 regulatory network on chlorophyll degradation and carotenoid biosynthesis pathways in citrus fruit. CrWRKY42 directly binds to the W-box element on the promoters of carotenoid biosynthesis genes (*CrPDS, CrLCYB2,* and *CrBCH1*) and chlorophyll degradation gene *CrSGR* to activate their expressions, thus promoting carotenoid biosynthesis and chlorophyll degradation, eventually accelerating the color conversion of citrus fruits. *CrSGR*, STAY-GREEN; *CrPDS*, phytoene desaturase; *CrLCYB2,* lycopene *β*-cyclase 2; *CrBCH1*, *β*-carotene hydroxylase 1.

### CrWRKY42 is a direct positive regulator of *CrBCH1*

The content and composition of carotenoids vary among citrus cultivars (Nisar et al., 2015). In general, what most mandarin cultivars primarily accumulate is *β-*cryptoxanthin and zeaxanthin (Ikoma et al., 2016). *CrBCH1* is an essential hydroxylase responsible for the hydroxylation of *β*-carotene into *β*-cryptoxanthin and zeaxanthin in citrus. However, the regulation of *CrBCH1* is still poorly understood. Fortunately, ‘Shiranuhi’ and its bud mutant ‘Jinlegan’ provided us with clues to study the transcriptional regulation of *CrBCH1* (Chen et al., 2023a). In the present study, we identified a WRKY transcription factor, CrWRKY42, and found that CrWRKY42 directly bound to W-box on the promoter of *CrBCH1* and activated its expression (Figure 3). In addition, we also found that the expression level of *CrBCH1* was significantly higher in both *CrWRKY42-*overexpression citrus calli and citrus fruits than in the control (Figure 4, 6), which provided additional evidence that CrWRKY42 induced the expression of *CrBCH1*. Additionally, we found a decrease in *β*-carotene content and a significant increase in zeaxanthin and violaxanthin content in *CrWRKY42-*overexpression citrus calli, which were complementary to the changes in carotenoid accumulation in the flavedo of ‘Jinlegan’, suggesting that CrWRKY42 might be a key gene contributing to the altered color of ‘Jinlegan’.

Furthermore, the studies related to the WRKY family in citrus have primarily focused on disease resistance. For instance, CsWRKY70 is involved in the establishment of resistance of citrus fruits to *Penicillium digitatum* (Deng et al., 2020). CsWRKY22 regulates canker susceptibility in sweet orange by enhancing cell expansion and *CsLOB1* expression (Long et al., 2021). CitWRKY28 promotes cuticle wax synthesis by activating the expression of *CitKCS* in citrus fruits (Yang et al., 2022). However, there are few studies on the regulation of carotenoid biosynthesis. This study provides new insights into the functions of the WRKY family in plants.

### CrWRKY42 regulates the *β* branch metabolism of carotenoids in citrus

Overexpression of *CrWRKY42* in citrus calli and citrus fruits significantly enhanced carotenoid accumulation (Figure 4, 6). Furthermore, the expressions of several carotenoid biosynthesis genes were up-regulated in *CrWRKY42*-overexpression citrus calli and fruits, which was consistent with the significantly increased total carotenoid content in *CrWRKY42*-overexpression lines. We also confirmed that CrWRKY42 could directly bind to the promoters of *CrPDS* and *CrLCYB2* to activate their expression (Figure 8), which synergistically regulated carotenoid accumulation. This result is in line with some previous reports that one transcription factor can interact with multiple carotenoid biosynthesis genes to participate in carotenoid accumulation, such as CsMADS6, CsERF61, and CsMYC2 (Lu et al., 2018; Zhu et al., 2021a; Yue et al., 2022), which might be attributed to a direct regulation or with a feedback mechanism. The *β* branch of carotenoid biosynthesis, as a main primary pathway of carotenoid accumulation in citrus, is more complex and diverse than the *α* branch. The metabolites of *β* branch serve as the precursor for the synthesis of various hormones and volatiles in citrus, such as abscisic acid (ABA) (Lu and Li, 2008), In *CrWRKY42*-overexpression citrus calli, we observed that carotenoids preferentially increased metabolic flow of the *β* branch, which predominantly promoted the hydroxylation of *β*-carotene into downstream zeaxanthin, meanwhile increasing violaxanthin, but no significant changes in the content of lutein in *α* branch. This result was similar to the accumulation of carotenoids during the ripening process of ‘Shiranuhi’ fruits. The preference of CrWRKY42 to regulate the *β* branch pathway might be due to the fact that CrWRKY42 could also be involved in the regulation of the ABA metabolic pathway. Previous studies have indicated that WRKY transcription factors affect phytohormones such as ABA (Rushton et al., 2012). In this study, elevated ABA content was also detected in *CrWRKY42*-overexpression citrus calli (Supplemental Figure S1), suggesting that CrWRKY42 might be involved in the regulation of the ABA metabolic pathway, but this needs to be further verified in future studies. In addition, the regulation of carotenoid metabolic pathways depends on different transcription factors. The overexpression of *CsMADS5* has been reported to principally affect the carotenoid content of the *α* branch, including the increased *α*-carotene and the decreased lutein content (Lu et al., 2021). CrMYB68, as a negative regulator, inhibits carotenoid synthesis in both *α* and *β* branch carotenoids (Zhu et al., 2017). SlWRKY35 enhances carotenoid accumulation (lutein, lycopene, and *β-*carotene) from upstream principally by directly binding to the promoter of *SlDXS1* and activating its expression (Yuan et al., 2022). In this study, we found that CrWRKY42 was critical for the targeted alteration of carotenoid content in the *β* branch in citrus.

### CrWRKY42 promotes color conversion of citrus fruits by accelerating chlorophyll degradation and carotenoid accumulation

The fruit ripening accompanies complex physiological processes, such as fruit softening, chlorophyll degradation, and carotenoid accumulation (Prasanna et al., 2007). In the present study, CrWRKY42 up-regulated the expression of *CrSGR* in *CrWRKY42*-overexpressed citrus fruits, but down-regulated the expression of *CrSGR* in *CrWRKY42*-RNAi fruits, suggesting that CrWRKY42 positively regulated the expression of *CrSGR*, thus participating in chlorophyll degradation. Our results further confirmed that CrWRKY42 directly bound to and activated the promoter of *CrSGR* (Figure 8). It is not uncommon that genes involved in different metabolic pathways are synergistically regulated by the identical transcription factor. In tomato, SlBBX20 is identified as a novel regulator that simultaneously enhances carotenoid and chlorophyll biosynthesis (Xiong et al., 2019). AdMYB7 regulates both carotenoid and chlorophyll biosynthesis by inducing the expression of key chlorophyll and carotenoid synthesis genes (Ampomah-Dwamena et al., 2019). In citrus, CsMADS3 coordinates *CsPSY1*, *CsLCYB2*, and *CsSGR* to regulate carotenoid biosynthesis and chlorophyll degradation (Zhu et al., 2023). In this study, CrWRKY42 accelerated chlorophyll degradation and carotenoid accumulation by up-regulating the expression of genes in both pathways, in turn promoting the degreening process and color conversion of fruits, which suggested that CrWRKY42 acted as a positive regulator of chlorophyll degradation and carotenoid accumulation to facilitate the coloration of citrus fruits (Figure. 6, 7).

In conclusion, our study revealed that CrWRKY42 synergistically regulated chlorophyll degradation and carotenoid biosynthesis by directly binding and activating genes related to both pathways, thus promoting the conversion of citrus fruit color. This study provides new insights into the regulation of chlorophyll degradation and carotenoid biosynthesis by the WRKY transcription factors during citrus fruit ripening.

## Materials and Methods

### Plant materials

The fruits of both the yellowish mutant ‘Jinlegan’ (MT) and its wild type ‘Shiranuhi’ (WT) tangor (*Citrus reticulata*) were sampled at different developmental stages (170, 190, and 210 days after flowering) in Chongqing, China. All the samples were frozen in liquid nitrogen after collection and stored at -80°C for further analysis. The citrus calli used for stable transformation were derived from Marsh grapefruit (*Citrus paradisi* Macf., ‘Rm’) and subcultured at 25℃ on solid Murashige Tucker (MT) medium. Fruits of ‘Valencia’ orange (*C. sinensis* L. Osbeck) were used for transient transformation. All the above materials were sampled with three biological replicates.

### RNA extraction, semi-quantitative PCR, and real-time quantitative PCR

The total RNA of samples was extracted using the kit (Aidlab Biotechnology, Beijing, China), following the instructions. cDNA was synthesized using HiScript II Q RT SuperMix for qPCR (+ gDNA wiper; Vazyme). Semi-quantitative PCR was performed using a PCR amplification program with 31 cycles. RT-qPCR was conducted using a Roche LightCycler 480 system (Roche) with three biological replicates. The RT-qPCR primers used in this study are listed in Supplemental Table S1.

### CrWRKY42 protein accession number and phylogenetic analysis

CrWRKY42 sequence data has been submitted to the GenBank database under accession number OR227423. Detailed sequence information of the genes in this study can be obtained in Phytozome (https://phytozome-next.jgi.doe.gov/) and NCBI database with the accession numbers listed in Supplementary Table S2. The phylogenic trees were constructed using RAxML-8.2.12 with the maximum likelihood method under PROTGAMMAAUTO model and 1,000 bootstrap replicates.

### Subcellular localization

The coding DNA sequence (CDS) of *CrWRKY42* without stop codon was cloned into PRI121 and transformed into *Agrobacterium tumefaciens* GV3101 for preparation. *Agrobacterium* cells carrying CrWRKY42-PRI121 and cytosolic marker H2B were mixed and co-transformed into leaves of tobacco (*Nicotiana benthamiana*) by transient injection. 3 days after injection, the fluorescence signal was observed and photographed using laser confocal microscopy (TCS SP8, Leica, Germany). The empty vector PRI121 was used as a negative control.

### Dual luciferase reporter assay

The dual luciferase reporter gene assay was performed by previously reported method with minor modifications (Lu et al., 2018). The effector vector pBD-CrWRKY42 was obtained by ligating the CDS sequence of *CrWRKY42* into the pBD vector. The empty pBD was used as a negative control, and the pBD-VP16 vector containing VP16 activation domain was used as a positive control. The pGreenII0800-LUC with five copies of GAL4 binding domain and an internal control REN driven by the 35S promoter was used as a reporter vector. The reporter vector and effector vector were transformed into *Agrobacterium tumefaciens* GV3101. Tobacco leaves were transiently injected with a mixture of effector and reporter at the ratio of 9:1, and the dual luciferase activity was measured 3 days after injection. The transcriptional activation activity of the gene was determined by the relative ratio of LUC/REN.

The full-length sequence of *CrWRKY42* was cloned into PK7WG2D, and the *CrBCH1*, *CrPDS*, *CrLCYB2*, and *CrSGR* promoter sequences were cloned into pGreenII0800-LUC. Tobacco leaves were injected with the mixture of effector and reporter at the ratio of 6:1, and the dual luciferase activity was measured 3 days after injection. The transcriptional activation activity of the gene was determined by the relative ratio of LUC/REN. LUC was imaged with the Plant Fluorescent Imaging System in *vivo* after the application of luciferin on the abaxial leaf surface. Three biological replicates were performed for each set of data.

### Transcriptional activation assay

The full-length sequence of *CrWRKY42* was cloned into pGBKT7, and the specific primer sequences were listed in Supplementary Table S1. Positive control (pGBKT7-53 + pGADT7-RecT), negative control (empty vector pGBKT7), and recombinant vector (pGBKT7-

CrWRKY42) were transformed into yeast (*Saccharomyces cerevisiae*) strain AH109, respectively. The transformed cells were incubated on SD/-Trp and SD/-Trp-His-Ade media for 3-7 days at 30°C. Transactivation activity was assessed by observing the growth of the transformed cells.

### Yeast one-hybrid (Y1H) assay

The full-length sequence of *CrWRKY42* was cloned into pGADT7 to construct the prey, and the promoter sequence of *CrBCH1* and *CrSGR* were cloned into the pAbAi vector to construct the bait. Y1H assay was performed by previously described method (Lu et al., 2018). **Electrophoretic mobility shift assay (EMSA)**

The CDS of *CrWRKY42* was cloned into PMAL-C_2_X to obtain the recombinant vector. The recombinant vector was transformed into *E. coli* BL21 (DE3) and then induced with 0.1 mmol/L IPTG at 16°C for 16 h, followed by the recombinant protein purification. 3’ Biotin-labeled probes were synthesized by Sangon company (Shanghai, China). The specific sequences were presented in Supplementary Table S1. MBP protein was used as a control, and unlabeled probes including identical or mutated oligonucleotides were used as cold competitors. EMSA was conducted by the previously reported method (Yue et al., 2020).

### Transformation of citrus calli

The full-length sequence of *CrWRKY42* was cloned into PH7WGAD for overexpression, and a specific fragment of *CrWRKY42* was inserted into the pHellsgate8-HG for interference. The specific primer sequences were listed in the Supplementary Table S1. The transformation was performed by previously described method (Lu et al., 2018).

### Carotenoid detection and analysis

Carotenoid extraction, detection, and analysis were performed, as described previously (Zhu et al., 2022). The samples were dissolved in ethyl acetate for high-performance liquid chromatography (HPLC, e2695; Waters, USA) analysis. The retention time and absorption spectra of samples were compared to those of standards to determine the carotenoids. Peak areas for phytoene, phytofluene, and other compounds were measured at 286, 348, and 450 nm, respectively. The carotenoid levels were quantified using calibration curves prepared with appropriate standards. The experiments were conducted with three biological replicates. The content of total carotenoid was measured by a UV-Visible spectrophotometer (UV-1700, Kyoto, Japan) at 450 nm by previously reported method (Lee and Castle, 2001).

### Citrus fruit transient transformation

The transient transformation of citrus fruits was performed according to the previous description (Gong et al., 2021). Briefly, the full-length or specific fragments of *CrWRKY42* were inserted into PH7WGAD or pK7GWIWG2D for overexpression or interference. The empty vector pH7WG2D or pK7GWIWG2D was used as a negative control. After infection, the fruits of *CrWRKY42-*overexpression and control (PK7-EV) were incubated in the storage chamber (temperature: 25℃), and the fruits of *CrWRKY42-*RNAi and control (RNAi-EV) were incubated in total darkness. At 10-18 days after incubation, the peel of the fruits near the injection site was sampled and frozen in liquid nitrogen for subsequent analysis. GFP fluorescence was used as a reporter to detect transgene expression in explants, which was observed with a fluorescent stereo microscope (SZX7; Mingmei, China). The citrus color index (CCI) [=1000×a /(L×b)] values were calculated. Three biological replicates were performed for each set of data.

### Total chlorophyll content detection and analysis

The extraction and detection of total chlorophyll in transgenic citrus fruits were performed as described in the previous report (Liu et al., 2004). Total chlorophyll is the sum of chlorophyll a and chlorophyll b.

### Detection and analysis of ABA content

The extraction and analysis of ABA content in the transgenic citrus calli have been described previously (Zhu et al., 2023).

### Statistical analysis

The data were analyzed using GraphPad 8.0. Student’s *t*-test and one-way ANOVA were conducted to determine significant differences between the two data sets at the levels of *P* < 0.05 and *P* < 0.01. The data were expressed as mean ± SD of at least three biological replicates.

## Supplemental Data

**Supplemental Figure S1. The ABA content in *CrWRKY42*-overexpression citrus calli.**

**Supplemental Table S1. Primers used in this study.**

**Supplemental Table S2. The accession numbers of related protein in this study.**

## Acknowledgments and Funding

We thank the Prof. Wang Pengwei (Huazhong Agricultural University, Wuhan, China) for providing the pHellsgate8-HG plasmid. This work was funded by the National Key Research and Development Program of China (No. 2022YFF1003100) and National Natural Science Foundation of China (No. 31930095).

## Author contributions

X.D. conceived and coordinated this project; H.C. and X.D. designed the experiments; H.C. performed the experiments with contributions from H.J., W.H., Z.Z., S.Z.; H.C. wrote the manuscript, under the supervision of and with contributions from X.D., J.Y., K.Z and L.C.

